# Stone Alkaline Water Induces Apoptosis of Prostate Cancer Cells and Inhibits Tumor Cell Induced Angiogenesis In Vitro

**DOI:** 10.1101/2019.12.15.877233

**Authors:** Sila Appak-Baskoy, lknur Kulcanay Sahin, Ozgun Teksoy, Mustafa Cengiz, Asuman Deveci Ozkan, Gamze Guney Eskiler, Namık Bilici, Pinar Oztopcu-Vatan, Yılmaz Altuner, Adnan Ayhanci

**Author notes:** Correspondence: Prof Adnan Ayhanci (PhD), Eskisehir Osmangazi University, Department of Biology, Meselik 26480 Eskisehir TURKEY, E mail.

## Abstract

Treatment options to improve overall survival rate of prostate cancer patients are limited since tumor cells acquire resistance to the chemotherapeutic drugs. We aimed to determine anticancer effects of stone alkaline water (SAW) on PC-3 and DU-145 prostate adenocarcinoma cell lines. SAW was obtained by triturating high stones under vacuum at 3000 °C. High mineral and trace element containing fraction of SAW was used for the experiments. Viability of the tumor cells was analyzed using tetrazolium based WST-1 cell proliferation assay, cell cycle analysis was carried out with Propidum Iodide staining (Muse™ Cell Cycle Kit). Acridine Orange and Annexin V stainings were done to analyze the cellular morphology and to determine apoptosis. Tumor cell derived angiogenesis was analyzed with migration and tube formation assays. SAW treatment resulted in accumulation of cells at G0/G1 phase and inhibited tumor cell induced HUVEC tube formation and migration. SAW treatment significantly decreased viability of PC-3 and DU-145 prostate adenocarcinoma cells and induced apoptotic cell death. Intriguingly, treatment of the prostate cancer cells with SAW inhibited tumor cell derived angiogenesis. SAW may aid in treating prostate cancer and molecules important for SAW’s apoptotic and anti-angiogenic effects need to be determined.

## Introduction

Prostate cancer is the second most common cancer, as well as, the second leading cause of mortality among males in the world [1]. Although studies on lifestyle factors that promote formation of the prostate cancer show promising results, the etiology of prostate cancer is still unclear as there is no specific carcinogenic substance found to be responsible of the disease formation [2]. Reports indicate that prostate cancer manifests itself as a result of genetic factors as well as lifestyle, diet, social and environmental factors [3,4,5]. Although surgery, radiotherapy and chemotherapy improve overall survival, most of the patients develop resistance against the chemotherapeutic drugs and tumor progresses with metastasis. The survival of patients with metastatic prostate cancer remains very low and new alternative therapeutic strategies are urgently needed to inhibit tumor progression.

Water is a liquid substance and contains various minerals and thus has significant role in human health. The chemical polarity of water and the ability to form hydrogen bonds are its most important characteristics. Stone alkaline water (SAW) is a high alkaline mineral water which is obtained by breaking certain stones under high heat, pressure and vacuum. The atomic absorption analysis (Table 1) shows that SAW contains variety of different minerals, mainly magnesium (Mg), calcium (Ca), sodium (Na) and carbonate (CO_3_).

**Table.1.** Mineral and trace element content of stone alkaline water (SAW)

| Content | Measured Values |
| --- | --- |
| Barium | 0,368 mg/L |
| Nickel | 0,012 mg/L |
| Magnesium | 4,25 mg/L |
| Iron | < 5 µg/L |
| Copper | < 1 µg/L |
| Chromium | < 2 µg/L |
| Sodium | 2,8 g/L |
| Sulfate | 0,25 g/L |
| Carbonate | 0,1 g/L |
| Quicksilver | Not found |
| Arsenic | Not found |
| Cyanide | Not found |
| Flor | Not found |
| Cadmium | Not found |
| Total solids | 375,5 g/L |
| Ph | 7,40 |
| Density | 1,2 g/cm<sup>3</sup> |

Mg and Ca are reported to protect cells against cancer [6,7]. Mg prevents DNA damage caused by different carcinogenic agents [8]. In some studies, researchers have demonstrated that the level of Ca and Mg in water has a preventive effect on colorectal and gastric cancers [9-11]. In the literature, alkaline water or mineral water has been shown to have beneficial effects on human health [12-16]. Nevertheless, there is a lack of evidence whether SAW and its mineral and trace element content has anti-cancer effects or not. Thus, the aim of this study is to examine the potential anti-cancer and apoptotic effects of SAW on metastatic prostate cancer cell lines PC-3 and DU-145 and how it affects tumor cell induced angiogenesis, *in vitro.*

## Material and Methods

### SAW

Stones with high carbonate content are washed and dried and then milled and sieved. The sieved stones were triturated under vacuum at 3000 °C after treating them with acid. A high mineral and trace element containing fraction of SAW was obtained by irrigating the stone powder a few hours. Gathered SAW was placed in transparent glass bottles and stored at room temperature.

### Cell Culture

PC-3 and DU-145 cells were purchased from the American Type Culture Collection (ATCC). Prostate tumor cells were incubated in Roswell Park Memorial Institute-1640 (RPMI-1640, Thermo) medium whereas HUVECs were maintained in DMEM medium with 10% heat-inactivated fetal bovine serum (FBS, Gibco) and 1% penicillin/streptomycin (Gibco). Cells were cultured in a humidified 37 °C incubator (Thermo Fisher Scientific) with 5% CO_2_. Both cells were counted using Countes II Cell Counter (Thermo Fisher Scientific, USA) for further analysis.

### WST-1 Analysis

In order to analyze the proliferation of the cells with/without SAW treatment tetrazolium based WST-1 cell proliferation assay was performed. Briefly, PC-3, DU-145 cells and HUVECs (2×10^4^ cells/well) were cultured in 96-well plates, then treated with different SAW concentrations (0.5-10 mg/mL) for 24 and 48 h. After incubation, cells were stained with 10 µL of WST-1 reagent (Biovision) at 37°C for 30 minutes in the dark and analyzed at 450 nm with 96-well plate reader (Allsheng, China). Each experiment was performed three times. We selected the most effective concentrations (1 and 2.5 mg/mL) of SAW and exposure time (24 h) for further experiments such as Annexin V, Acridine Orange stainings and cell cycle analysis.

### Annexin V Analysis

In order to show the possible apoptotic effect of SAW, Annexin V analysis was carried out. For this purpose, PC-3, DU-145 cells and HUVECs (1×10^5^ cells/well) were cultured in six-well plates for 24 h and treated with 1 and 2.5 mg/mL of SAW for 24 h with fresh medium. After treatment, the cells were washed twice with cold phosphate buffer (PBS) after centrifugation at 2000x g for 5 minutes. Cells were stained with Muse™ Annexin V & Dead Cell Assay kit (Millipore) and incubated for 30 minutes at room temperature in dark conditions. At the end of incubation cells were analyzed using the Muse™ Cell Analyzer (Millipore). Each experiment was performed in three replicates for each of the cell line.

### Cell Cycle Analysis

For the cell cycle analysis, Propidum Iodide based nuclear staining was carried out according to the manufacturer’s protocol. Briefly, PC-3, DU-145 cells and HUVECs (5×10^5^ cells/well) were cultured in six-well plates for 24 h and treated with 1 and 2.5 mg/mL of SAW for 24 h with fresh medium. After the treatment, cells were fixed in 70% ethyl alcohol (EtOH) and stored at -20 °C for at least three hours. Fixed cells were washed with cold PBS and centrifuged at 2000 x g for 5 minutes and stained using a Muse™ Cell Cycle Kit (Merck Millipore) for 30 minutes in dark conditions. After the staining, analysis was done by Muse™ Cell Analyzer (Merck Millipore). Each experiment was performed in three replicates for each cell line.

### Acridine Orange Staining

Cellular morphology of PC-3, DU-145 cells and HUVECs was analyzed with Acridine Orange staining. For this purpose, 5×10^5^ cells/well were cultured in six-well plates for 24 h and treated with 1 and 2.5 mg/mL of SAW for 24 h. After incubation, cells were fixed and stained by adding 500 µL of acridine orange for 30 minutes in the dark and examined with EVOS FL Cell Imaging System (Thermo).

### *In vitro* Angiogenesis Assays

PC-3 and DU-145 cells that were treated with SAW and untreated control cells were replaced with starvation medium (RPMI without FBS). Cells were starved overnight and supernatants were collected and used for stimulation of HUVECs in tube formation (2D angiogenesis) and migration assays.

The tube formation assay was performed on a 48-well tissue culture dish that was coated with 150 µl growth factor-reduced Matrigel (BD) that was left to solidify at 37°C for 30 min. 25,000 HUVECs were resuspended in the subsequent starvation media of the tumor cells and seeded on Matrigel-coated wells. Bright field images of capillary-like tubes were taken after 8h of incubation at 37°C. The mean branch points were quantified using Image J.

Boyden chamber migration assay was run on transwell inserts (5 µm pore size- Corning) that were coated with 0.1% gelatin for 10 minutes at 37 °C then washed with PBS. HUVECs were suspended in media supplemented with 0.1% FBS and were seeded onto the transwell inserts. Migration of HUVECs was stimulated with the control and SAW treated cells’ starvation media in the lower well. HUVECs were allowed to migrate for 16h. At the end of the assay, supernatants were aspirated, transwell inserts were washed with PBS and fixed with ice-cold methanol. HUVECs were stained with Hoechst dye to visualize the migrated cells. Nuclei were quantified using Image J.

### Statistical Analysis

All data was statistically evaluated by SPSS 22.0. Data were expressed as a mean ± SD of three experiments. The Analysis of Variance (ANOVA) test by Tukey’s test was conducted for multiple comparison. p<0.05 was considered to be statistically significant.

## Results

### The Cytotoxic Effects of SAW

WST-1 analysis was performed to evaluate the cytotoxic effects of SAW on PC-3, DU-145 cells and HUVECs (Figure 1). Viability of PC-3 and DU-145 cells were decreased in a dose-dependent manner. When PC-3 and DU-145 cells were treated with 1 mg/mL of SAW, the viability of cells significantly reduced to 51.44%, 58.55%, respectively for 24 h (p<0.05). However, 69.19% and 61.88% inhibition were found in PC-3 and DU-145 cells, respectively at 2.5 mg/mL for 24 h. The proliferation of HUVECs was reduced to 84.0% and 72.27% at 1 and 2.5 mg/mL of SAW for 24 h. Therefore, SAW treatment was found to be much more effective in inhibiting viability of PC-3 cells than DU-145 cells and it didn’t affect viability of HUVECs much in 24 h. Viability experiment revealed that, the most effective doses of SAW were 1 and 2.5 mg/mL administrated for 24 h, therefore these doses were selected for further analysis.

**Figure 1.**
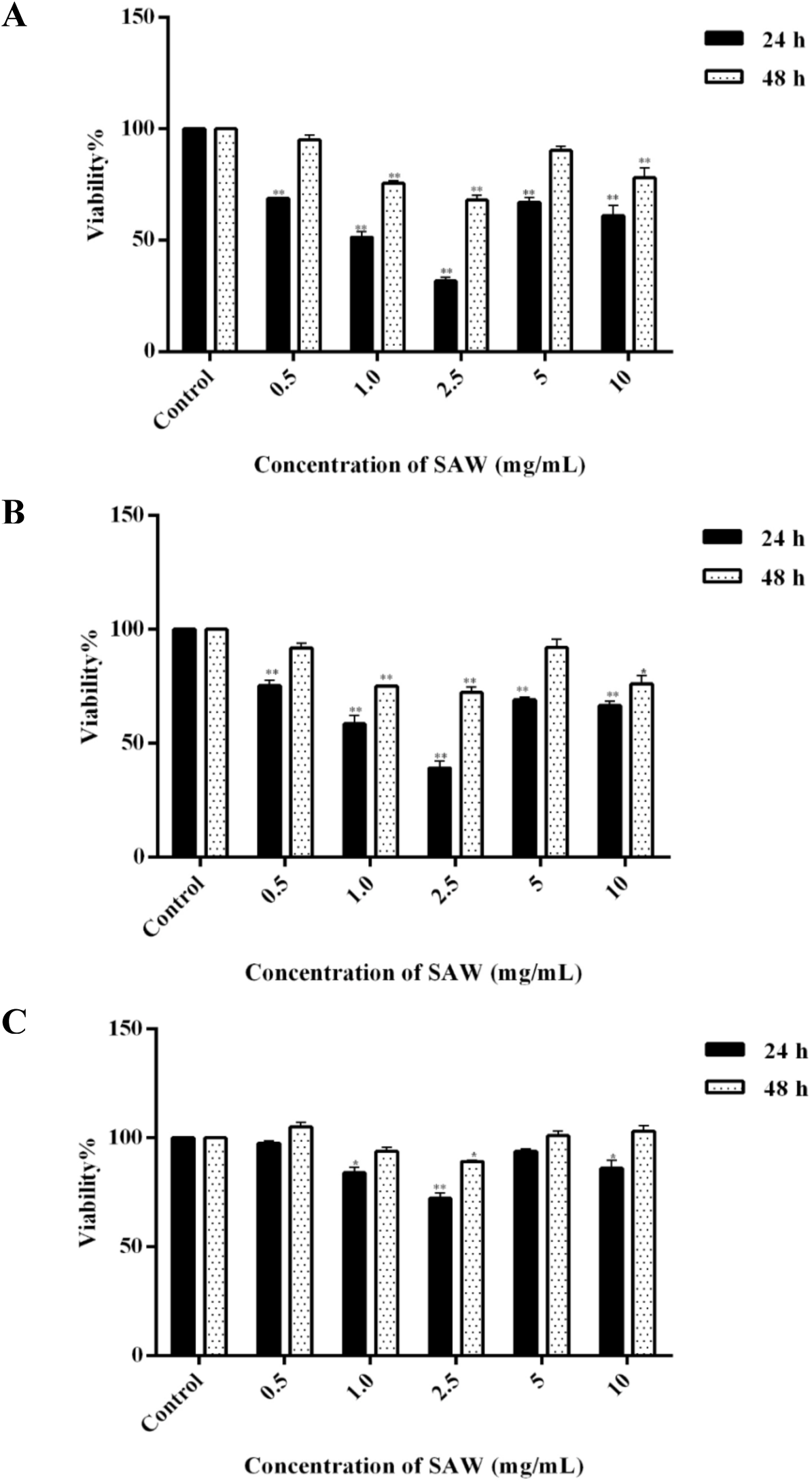
The viability of (A) PC-3, (B) DU-145 and (C) HUVEC cells exposed to 0.5-10 mg of SAW for 24 and 48 hours. (p<0.05*, p<0.01**).

### Apoptotic Effects of SAW

Annexin V analysis was conducted in order to determine the apoptotic effects of SAW on PC-3, DU-145 cells and HUVECs (Figure 2). Particularly, SAW significantly induced apoptosis in PC-3 and DU-145 cells after treatment of SAW (1 and 2.5 mg/mL) (p<0.05). The percentage of total apoptotic cells was remarkably increased to 77.65% and 66.25% in PC-3 and DU-145 cells, respectively at 2.5 mg/mL of SAW for 24 h compared to the control group (p<0.05). Furthermore, the proportion of total apoptotic cells was 19.69% and 26.31% at 1 and 2.5 mg/mL of SAW, respectively in HUVECs. The obtained results show that SAW caused apoptotic cell death and these findings were consistent with the results of WST-1 analysis.

**Figure 2.**
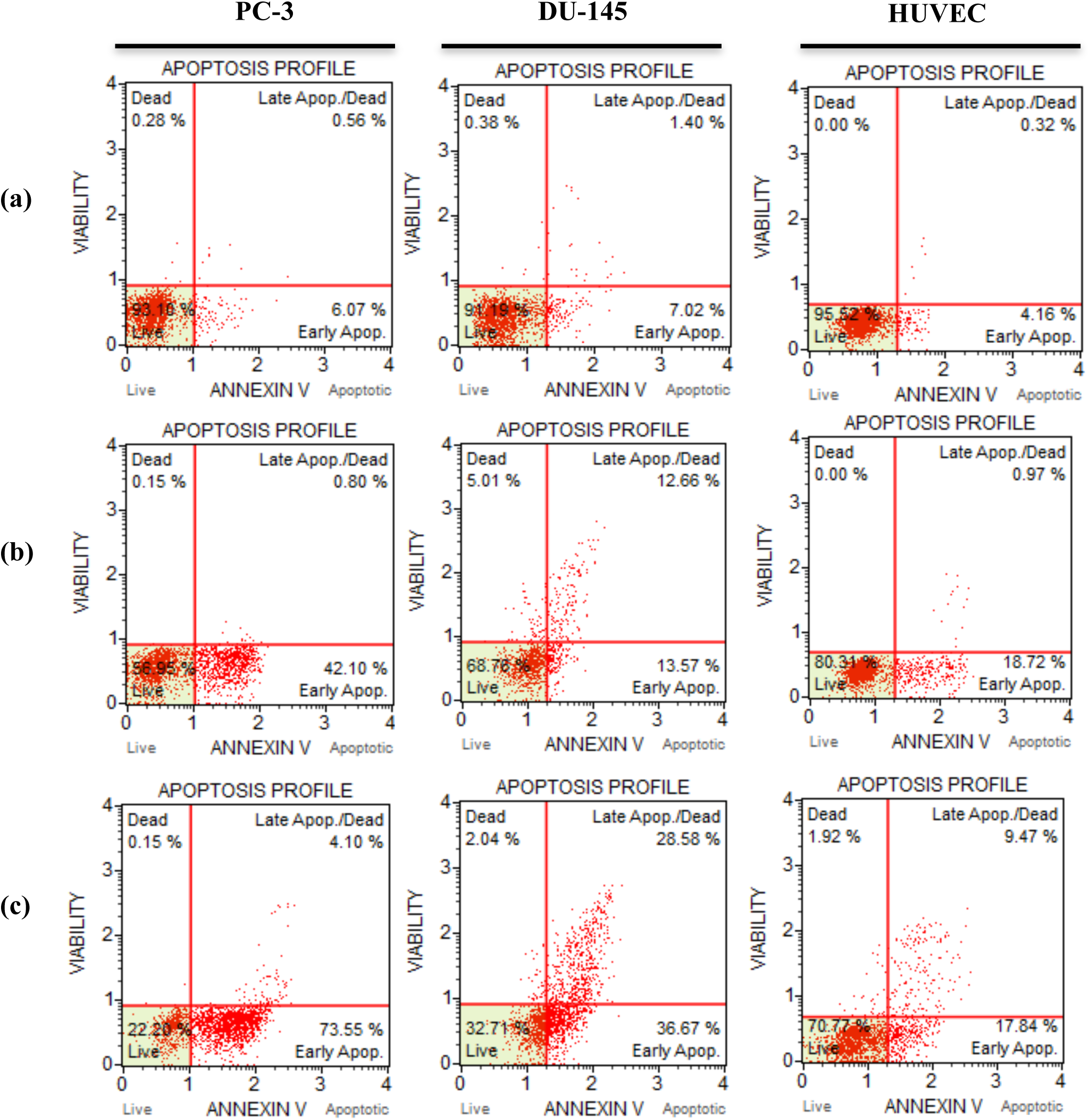
Annexin V analysis. of PC-3, DU-145 and HUVECs exposed to (b)1 mg/mL and (c) 2.5 mg/mL of SAW for 24 hours compared with (a) control.

### Cell Cycle Arrest

Results of the **c**ell cycle arrest analyses indicated that SAW treatment (1 and 2.5 mg/mL) resulted in G0/G1 arrest in PC-3, DU-145 and HUVECs (Figure 3). The percentage of PC-3, DU-145 cells and HUVECs in the G0/G1 phase were considerably increased to 92.6%, 89.5% and 75.3% at 2.5 mg/mL SAW concentrations compared to the control group (63.0%, 65.2% and 63.4%), respectively (p<0.05) (Figure 3A and 3B). Therefore, our findings showed that SAW treatment significantly accumulated cells in the G0/G1 phase arrest and induced apoptotic cell death.

**Figure 3.**
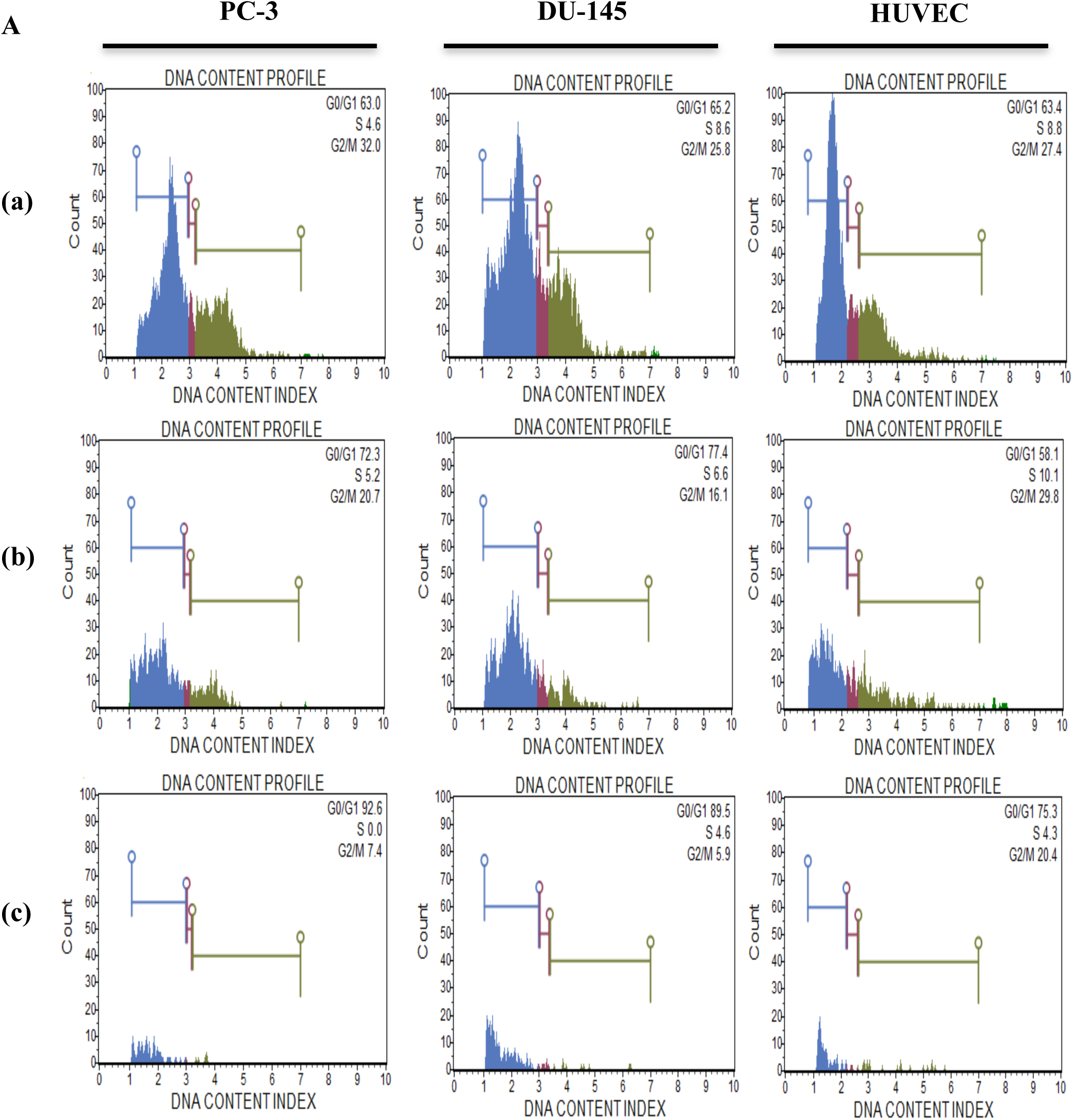
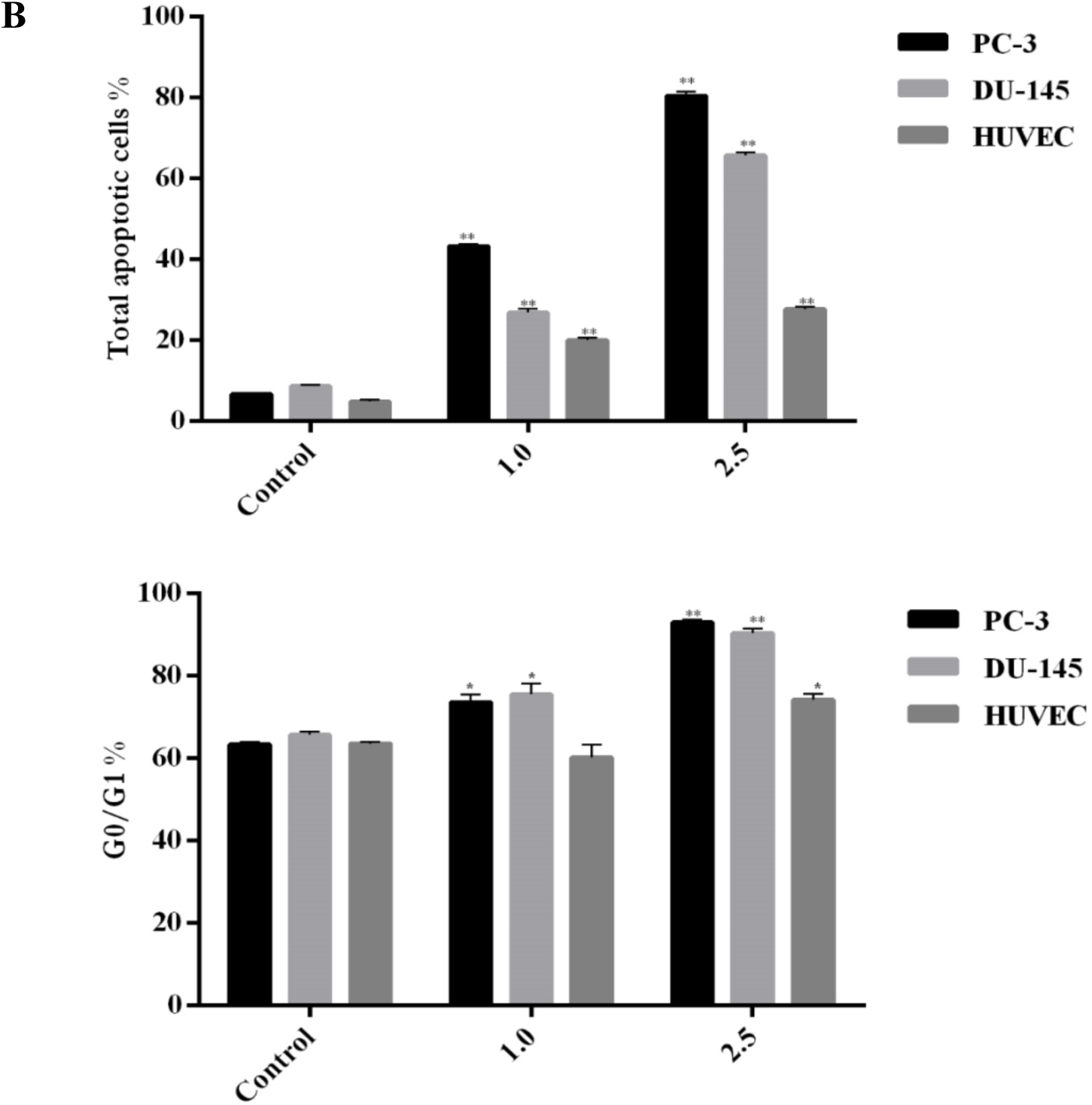
(A) Cell cycle analysis of PC-3, DU-145 and HUVECs following incubation with (b) 1 and (c) 2.5 mg/mL of SAW for 24 hours compared with control (a). (B) total apoptotic cell amount and accumulation of G0/G1 arrest in response to SAW in PC-3, DU-145 and HUVEC cells (p<0.05*, p<0.01**).

### Cell Morphology

The changes in the morphology of PC-3, DU-145 and HUVECs after treatment of SAW were determined by acridine orange staining (Figure 4). Apoptotic bodies and chromatin condensation were observed in PC-3 and DU-145 cells and some of the cells indicated necrotic morphology in response to SAW treatments. Cytoplasmic shrinkage and more round shape were only observed at 2.5 mg/mL concentration of SAW in PC-3 and DU-145 cells. Additionally, there were no specific morphological changes observed in HUVECs following incubation with SAW. However, there were some apoptotic bodies and necrotic cells. Consequently, PC-3 cells were more sensitive to SAW treatment than DU-145 cells and SAW treatment didn’t affect HUVECs morphology.

**Figure 4.**
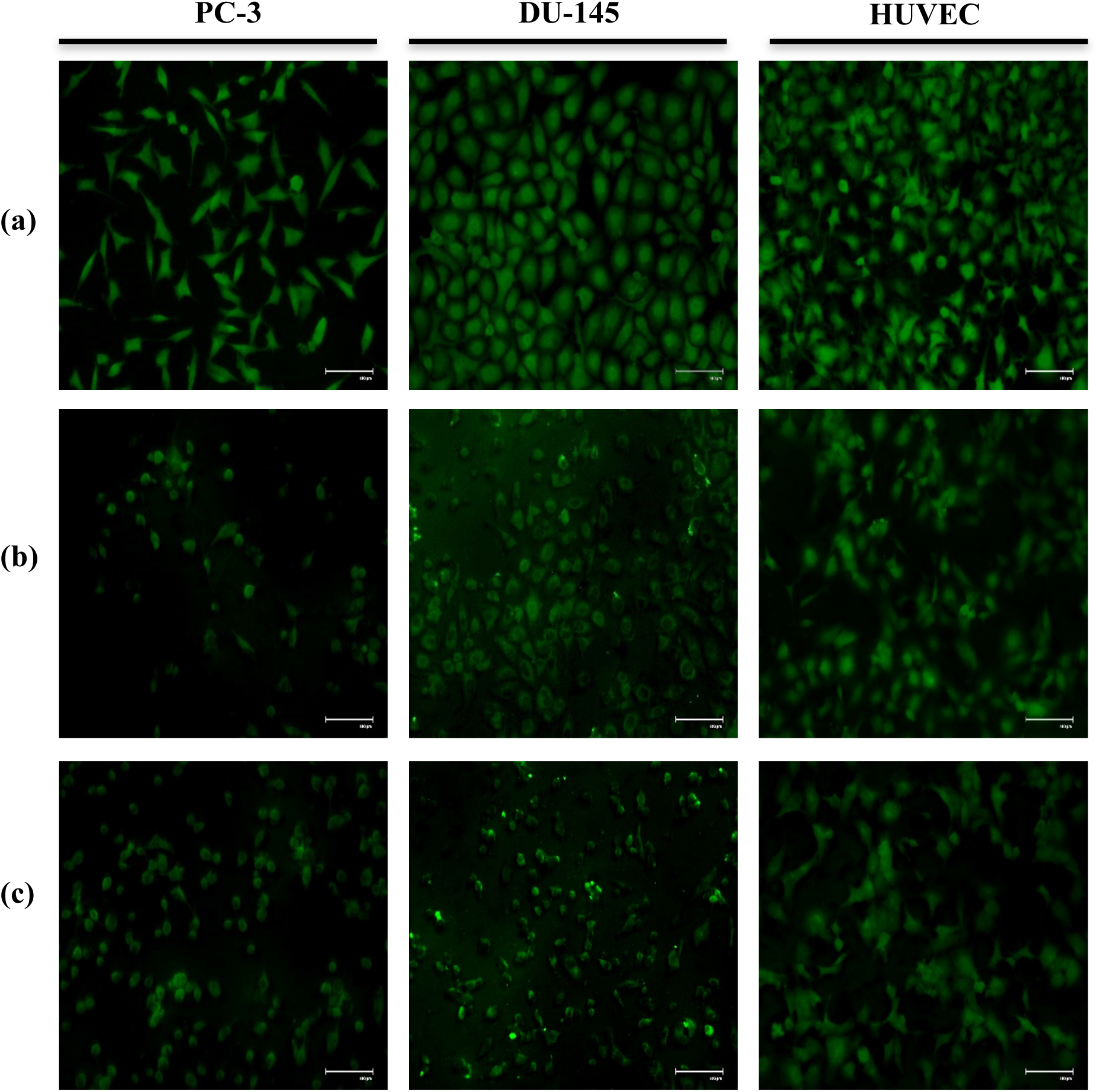
Acridine Orange staining of PC-3, DU-145 and HUVECs: a) control b) 1 mg/mL and c) 2.5 mg/mL SAW for 24 hours. Scale bar: 100 µm

### Inhibition of Tumor Cell Induced Angiogenesis

Boyden chamber migration assay showed that there were significantly less migrated HUVECs towards the supernatants of SAW treated PC-3 and DU-145 cells compared to control cells’ (Figure 5A-B). Moreover, 2D angiogenesis assay revealed that supernatants of SAW treated PC-3 and DU-145 significantly inhibited number of branch points HUVECs made on the matrigel matrix compared to control cells’ supernatants (Figure 5C-D). Taken together, these two experiments show that tumor cell secreted angiogenic factor(s) that induce angiogenesis was blocked by the SAW treatment.

**Figure 5:**
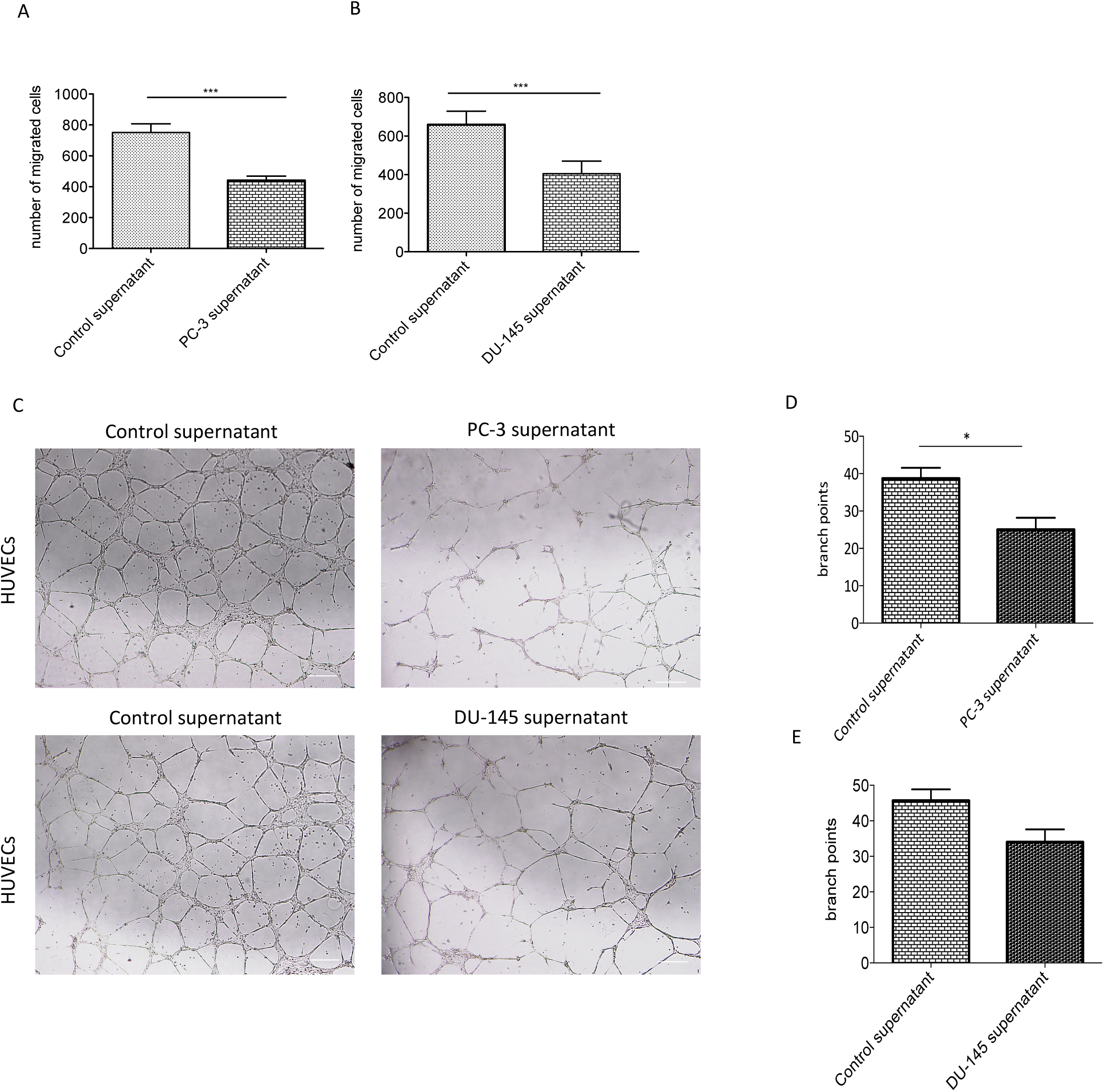
HUVECs functional assays with conditioned medium of SAW treated PC-3 and DU-145 cells. A. Migration of HUVECs stimulated with control and SAW treated PC-3 cells B. Migration of HUVECs stimulated with control and SAW treated DU-145 cells C. Light microscope pictures of HUVECs forming tubes in 2D angiogenesis assay stimulated with control and SAW treated PC-3 and DU-145 cells D. Number of branchpoints HUVECs make with conditioned medium of control and SAW treated PC-3 cells E. Number of branchpoints HUVECs make with conditioned medium of control and SAW treated Du-145 cells (p<0.05*, p<0.01**), Scale bar: 50 µm

## Discussion

In the current study, SAW’s anti-cancer effects were investigated on metastatic prostate cancer cell lines, for the first time. Our findings demonstrated that SAW induced apoptotic cell death and cell cycle arrest and suppressed cell viability in metastatic prostate cancer cells and their ability to induce angiogenesis, *in vitro*.

SAW is a high alkaline mineral water which contains different minerals including Na and Mg and may have anti-cancer effects due to its rich mineral content. Na and Mg, the most abundant minerals in the body, have been shown to regulate glucose metabolism, inflammation, and cell proliferation [16]. Thus Na and Mg are involved in a large number of biological processes, so the lack of these minerals can affect many processes including tumor progression. Low concentration of Mg is associated with an increase risk of various diseases such as cancer [17]. For example, intake of magnesium has been reported to be protective against colorectal cancer progression [18].

PC-3 and DU-145 cell lines are both androgen receptor (AR) positive prostate adenocarcinoma cell lines. PC-3 cells have more metastatic potential than DU-145 cells. PC-3 cells are obtained from bone metastasized prostate cancer whereas DU-145 cells are derived from brain metastasis and these cells do not express prostate specific antigen [19]. These two prostate cell lines have different genetic backrounds. PC-3 cells are found to carry deletion at codon 138 in TP53 gene associated with loss of a 17p (hemizygous 17p) whereas DU-145 cells have two mutations in TP53 gene [20]. Thus, genetic background and origin may cause differences in responses of PC-3 and DU-145 cells against therapeutic agents and this may also be the reason why two cell lines respond differently to SAW.

Growing tumor mass needs oxygen and nutrients and vasculature surrounding the tumor is not sufficient to replenish the tumor cells after tumor reaches a size of approximately 1 mm^3^. In order to make their own vasculature tumor cells secrete angiogenic factors that derive the formation of new tumor vasculature from the existing vessels of the organ. Tumor cells not only use new vasculature to get oxygen and nutrients but they can also use this newly formed vasculature as a route for metastatic dissemination. Tumor cells in vitro also secrete these angiogenic factors especially when the cells are nutrient deprived. Therefore, we used the starvation medium of SAW treated PC-3 and DU-145 cells to stimulate HUVECs in migration and tube formation experiments and found out that SAW treatment of tumor cells significantly inhibits tumor induced angiogenesis *in vitro*. One of the main molecules that derive angiogenesis is VEGF-A that is triggered by hypoxia and it is expressed by both PC-3 and DU-145 cells [21]. SAW may inhibit angiogenesis by inhibiting release of this growth factor by the tumor cells.

In conclusion, our findings showed that SAW had a significant suppressive effect on cell proliferation, tumor derived angiogenesis and it induced apoptotic cell death in prostate cancer cells. However, the molecules responsible for its apoptotic effects and the inhibition of tumor derived angiogenesis need to be further investigated mechanistically. Thus, SAW might be potentially promising therapy for treatment of metastatic prostate cancer.

## Acknowledgements

We thank Mr. Faruk Durukan for the preparation of SAW.

